# Asymmetry Map of Human Brain in Young Adults

**DOI:** 10.1101/2020.12.03.409805

**Authors:** Maryam Malekzadeh, Alireza Kashani

## Abstract

Although, asymmetry is a central organizational aspect of human brain, it has not been clearly described yet. Here, we have studied structural brain asymmetry in 1113 young adults using data obtained from Human Connectome Project. A significant rightward asymmetry in mean global cerebral cortical thickness, surface area and gray matter volume as well as volumes of cerebral white matter, cerebellar cortex and white matter, hippocampus, putamen, caudate nucleus, nucleus accumbens and amygdala was observed. Thalamus showed a leftward asymmetry. Regionally, most cerebral cortical regions show a significant rightward asymmetry in thickness. However, cortical surface area and gray matter volume are more evenly distributed between two hemispheres with almost half of the regions showing a leftward asymmetry. In addition, a strong correlation between cortical surface area and gray matter volume as well as their asymmetry indices was noted which results in concordant asymmetry patterns between cortical surface area and gray matter volume in most cortical regions.

## INTRODUCTION

Human brain is a lateralized structure (1). Several lines of evidence suggest that this feature is centrally involved in numerous tasks including cognitive and affective processes in physiological as well as pathological states (2–7). However, despite over a century of research, this important hallmark has not been clearly understood yet. To date, different and frequently conflicting reports about how human brain is lateralized and affected by variables like gender and age have been published (8–14). Variability in methods and populations, subtle and varying nature of asymmetry with age as well as vast inherent individual variability are considered some of the reasons for previous inconsistent reports (11, 13, 15–18). Young Adult Human Connectome Project (HCP) has applied standardized parcellation and imaging methods to study 300 families of 22-35 years old twin and non-twin siblings which are diverse enough to reflects ethnic diversity of the United States (19). This provides an excellent opportunity to study brain asymmetry in young adults and how is affected by specific variables like gender. Furthermore, HCP data set allows us to study asymmetry in different cortical and subcortical structures in same group of participants since ever growing data indicate that they actively participate in cardinal brain functions (20–22). In this study, we have studied asymmetry patterns in three measures of the cerebral cortex: thickness, surface area, gray matter volume, as well as cerebral white matter, cerebellar cortex and white matter, hippocampus, pallidum, putamen, caudate nucleus, nucleus accumbens, amygdala and thalamus.

## MATERIALS AND METHODS

All analyses were performed on datasets downloaded from the Young Adult HCP database (19). The process of acquisition and processing of the MRI images are explained here (23). The structural data of 1113 participants, 606 female and 507 male, who were between 22-35 years old was analyzed. Statistical analyses were performed using SPSS software ver. 24.0 (IBM, Armonk, NY, USA). Asymmetry Index (AI) of thickness, surface area and gray matter volume of each cortical region as well as volumes of cerebral white matter, hippocampus, pallidum, putamen, caudate nucleus, nucleus accumbens, amygdala and thalamus was calculated for each subject according to the formula of 2 × (left-right)/(left+right). Positive and negative AIs indicate leftward or rightward asymmetry respectively. A single group *t-*test was carried out for comparison of AI means. A significance level of *P* < 0.0011 (0.05/44 brain regions) was assumed due to Bonferroni corrections to prevent inflated error rates. Effect size was expressed as Cohen’s-d (*d*). A Multivariate Analysis of Variance (MANOVA) test was conducted to evaluate effect of gender as between-subject factor and age as a covariate on AIs of studied structures. Results were analyzed with region specific *post hoc* tests. Coefficient of Variation (CV) was calculated according to the formula: CV= standard deviation/mean to evaluate variability of metrics in the cerebral cortex. Pearson correlation coefficient (R) was calculated to study relationship between them. For correlation analysis statistical significance threshold was set at *P* < 0.05.

## RESULTS

### Cerebral Cortex

A significant rightward asymmetry is observed in mean global cortical thickness, surface area as well as mean global cortical gray matter and cerebral white matter volumes. (Table.1 and Fig.1). However, at regional level differential asymmetry patterns were observed as follows.

**Table 1.** Global asymmetry index for each metric of cerebral cortex and white matter.

|  |  |  | AI |  |  | *P* |
| --- | --- | --- | --- | --- | --- | --- |
|  |  |  | Mean | SD | *d* |  |
| Cerebrum | Cortex | Thickness | -0.014 | 0.021 | -0.65 | 0.000000 |
|  |  | Surface Area | -0.006 | 0.011 | -0.54 | 0.000000 |
|  |  | Gray Volume | -0.018 | 0.019 | -0.98 | 0.000000 |
|  | White Matter |  | -0.014 | 0.009 | -1.51 | 0.000000 |

**Fig. 1.**
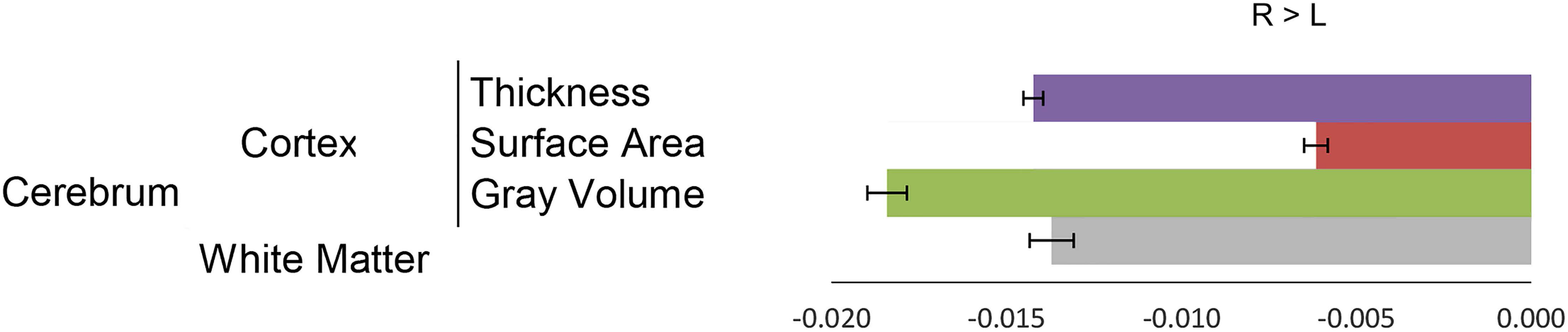
Global asymmetry indices for metrics of cerebral cortex and white mater volume. Negative asymmetry (R > L) indicates rightward asymmetry. Error bars represent SEM

#### 1. Thickness

85% (29/34) of cerebral cortical regions show a significant degree of asymmetry in thickness with 76% (26/34) presenting a rightward asymmetry. Insula, caudal anterior and posterior cingulate cortices show a significant leftward asymmetry. Parahippocampal and precentral gyri as well as rostral anterior cingulate, pericalcarine and cuneus cortices display a non-significant difference. All other regions show a significant rightward asymmetry in thickness (Table.2 and Fig.2).

**Table 2.** Asymmetry Index for thickness of each cortical region

| Region | AI |  |  | *P* |
| --- | --- | --- | --- | --- |
|  | Mean | SD | *d* |  |
| Caudal anterior cingulate cortex | 0.07 | 0.11 | 0.62 | 0.000000 |
| Insula | 0.01 | 0.05 | 0.28 | 0.000000 |
| Posterior cingulate gyrus | 0.01 | 0.07 | 0.21 | 0.000000 |
| Parahippocampal gyrus | 0.01 | 0.08 | 0.08 | 0.009564 |
| Rostral anterior cingulate cortex | 0.00 | 0.07 | 0.06 | 0.042102 |
| Precentral gyrus | 0.00 | 0.03 | 0.04 | 0.204130 |
| Pericalcarine cortex | 0.00 | 0.05 | -0.03 | 0.303877 |
| Cuneus cortex | 0.00 | 0.05 | -0.07 | 0.019816 |
| Frontal pole | -0.01 | 0.07 | -0.12 | 0.000041 |
| Lingual gyrus | -0.01 | 0.05 | -0.16 | 0.000000 |
| Isthmus of cingulate gyrus | -0.03 | 0.17 | -0.16 | 0.000000 |
| Lateral orbitofrontal cortex | -0.01 | 0.04 | -0.18 | 0.000000 |
| Fusiform gyrus | -0.01 | 0.03 | -0.19 | 0.000000 |
| Caudal middle frontal gyrus | -0.01 | 0.03 | -0.21 | 0.000000 |
| Rostral middle frontal gyrus | -0.01 | 0.03 | -0.22 | 0.000000 |
| Paracentral sulcus | -0.01 | 0.04 | -0.27 | 0.000000 |
| Postcentral gyrus | -0.01 | 0.03 | -0.32 | 0.000000 |
| Pars orbitalis | -0.02 | 0.05 | -0.34 | 0.000000 |
| Inferior temporal gyrus | -0.01 | 0.04 | -0.35 | 0.000000 |
| Pars opercularis | -0.02 | 0.04 | -0.40 | 0.000000 |
| Precuneus cortex | -0.01 | 0.03 | -0.42 | 0.000000 |
| Superior frontal gyrus | -0.01 | 0.03 | -0.44 | 0.000000 |
| Entorhinal cortex | -0.04 | 0.08 | -0.47 | 0.000000 |
| Transverse temporal cortex | -0.03 | 0.06 | -0.47 | 0.000000 |
| Superior temporal gyrus | -0.02 | 0.03 | -0.57 | 0.000000 |
| Supramarginal gyrus | -0.02 | 0.03 | -0.58 | 0.000000 |
| Pars triangularis | -0.02 | 0.04 | -0.60 | 0.000000 |
| Lateral occipital cortex | -0.02 | 0.04 | -0.63 | 0.000000 |
| Superior parietal cortex | -0.02 | 0.03 | -0.64 | 0.000000 |
| Middle temporal gyrus | -0.02 | 0.03 | -0.73 | 0.000000 |
| Temporal pole | -0.06 | 0.08 | -0.80 | 0.000000 |
| Banks of superior temporal cortex | -0.04 | 0.05 | -0.82 | 0.000000 |
| Inferior parietal cortex | -0.03 | 0.03 | -0.95 | 0.000000 |
| Medial orbitofrontal cortex | -0.07 | 0.06 | -1.15 | 0.000000 |

**Fig. 2.**
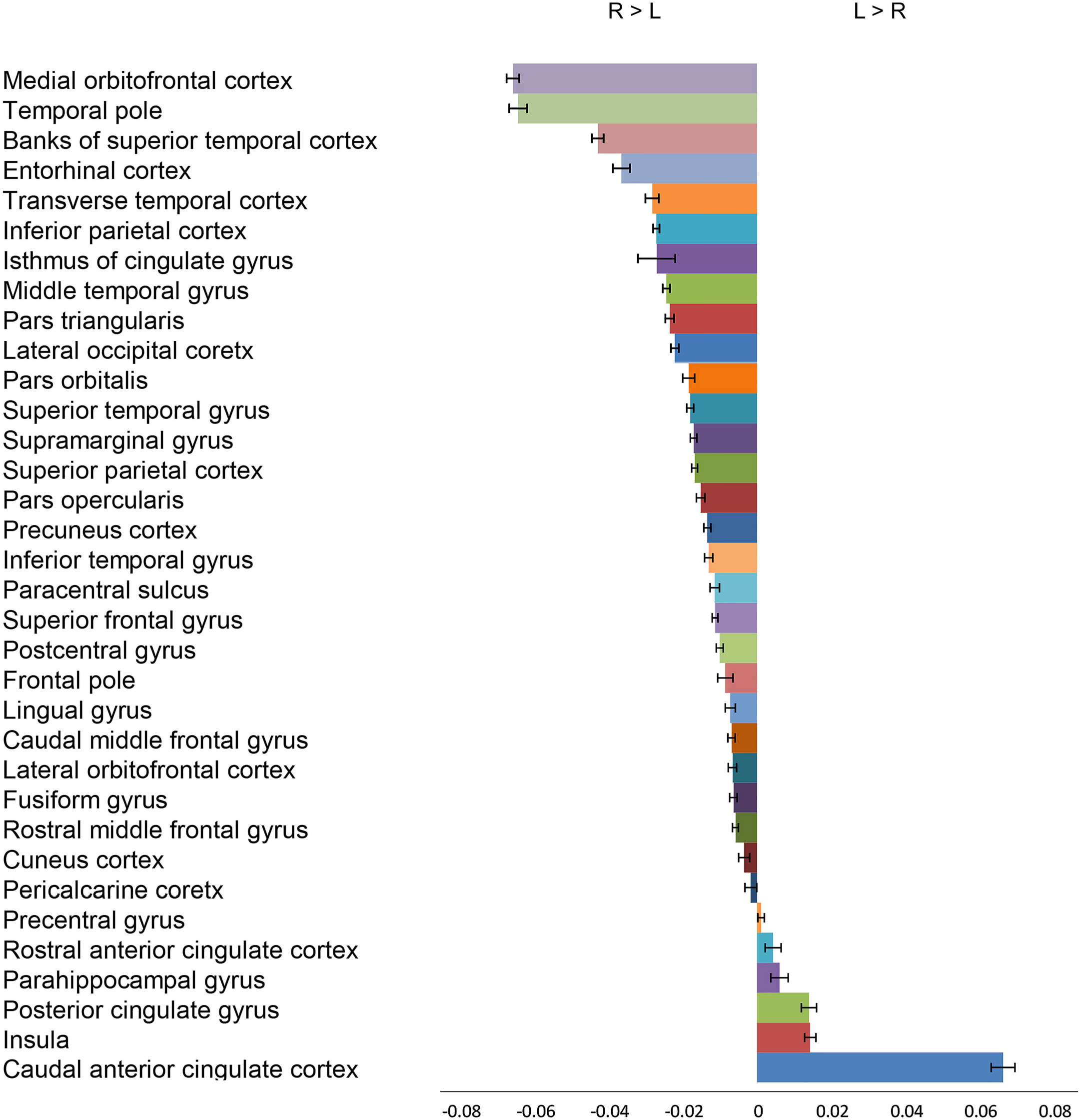
Asymmetry indices of thickness in cortical regions. Positive asymmetry indicates leftward asymmetry (L > R), while negative asymmetry (R > L) indicates rightward asymmetry. Error bars represent SEM.

#### 2. Area

94% (32/34) of the cerebral cortical regions present a significant asymmetry in surface area. However, distribution of regions between two hemispheres is more even relative to cortical thickness. 41% (14/34) show a rightward and 53% (18/34) present a leftward asymmetry. Regions with rightward asymmetry are widely distributed on all cerebral surfaces including frontal pole, pars orbitalis, pars triangularis, insula, middle temporal gyrus and inferior parietal, pericalcarine, caudal anterior cingulate cortices. Regions with leftward asymmetry show a tendency to peri-Sylvian areas and include superior temporal and supramarginal gyri, temporal pole, pars opercularis as well as transverse temporal, entorhinal and rostral anterior cingulate cortices (Table.3 and Fig.3).

**Table 3.**
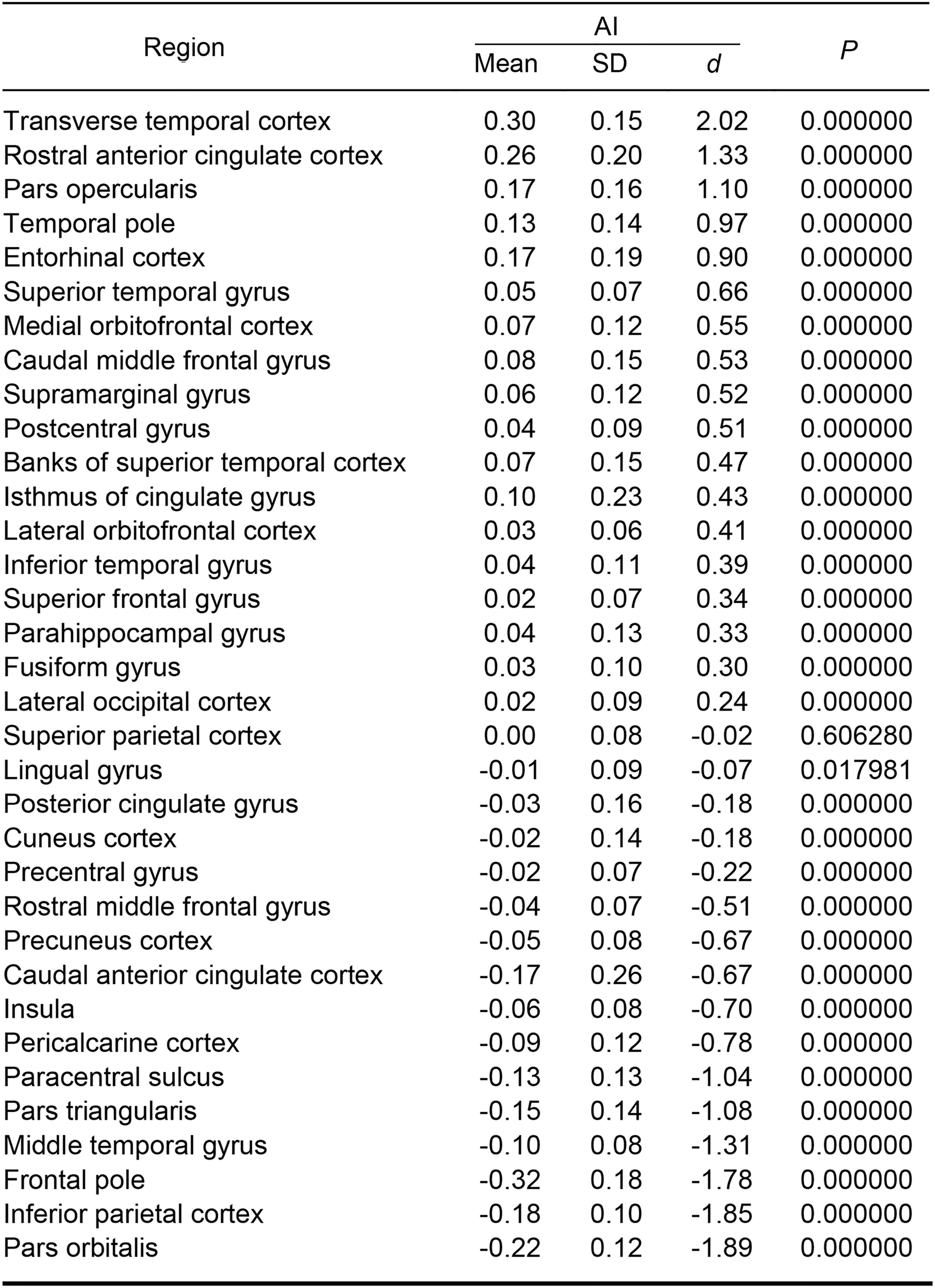
Asymmetry Index for surface area of each cortical region

**Fig. 3.**
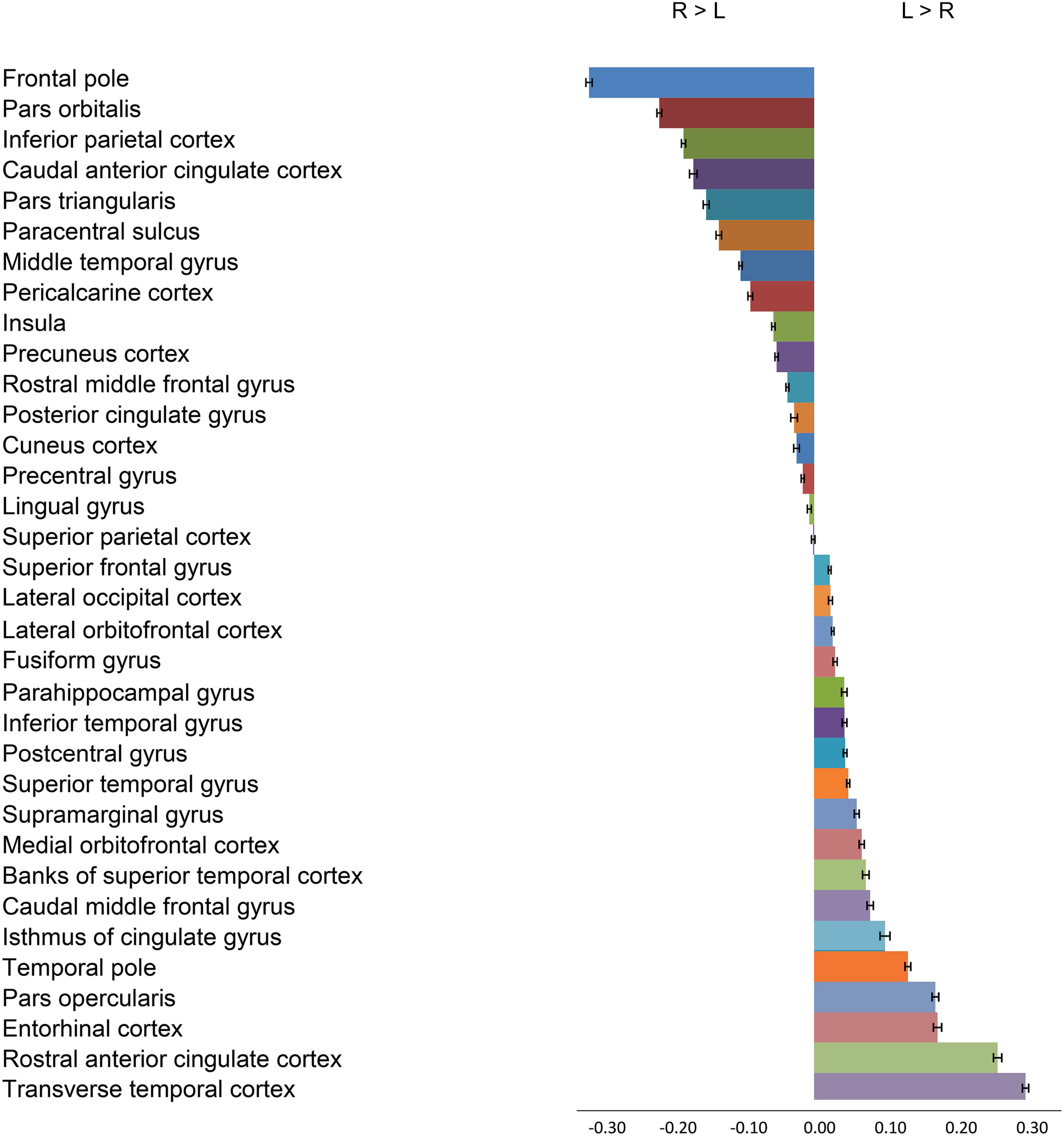
Asymmetry indices of surface area in cortical regions. Positive asymmetry indicates leftward asymmetry (L > R), while negative asymmetry (R > L) indicates rightward asymmetry. Error bars represent SEM.

#### 3. Gray matter volume

88% (30/34) of cerebral cortical regions show a significant asymmetry in their gray matter volumes. 41% (14/34) display a rightward and 47% (16/34) a leftward asymmetry. Regions with leftward asymmetry indices include transverse temporal, rostral anterior cingulate, pars opercularis, caudal anterior cingulate and entorhinal cortices. On the other hand, middle temporal gyrus, pars triangularis, banks of superior temporal sulcus, inferior parietal cortex, pars orbitalis and frontal pole regions present a rightward asymmetry (Table.4 and Fig.4).

**Table 4.** Asymmetry Index for gray matter volume of each cortical region

| Region | AI |  |  | *P* |
| --- | --- | --- | --- | --- |
|  | Mean | SD | *d* |  |
| Transverse temporal cortex | 0.25 | 0.17 | 1.45 | 0.000000 |
| Rostral anterior cingulate cortex | 0.25 | 0.22 | 1.15 | 0.000000 |
| Pars opercularis | 0.17 | 0.16 | 1.07 | 0.000000 |
| Caudal anterior cingulate cortex | 0.15 | 0.25 | 0.59 | 0.000000 |
| Entorhinal cortex | 0.11 | 0.19 | 0.59 | 0.000000 |
| Parahippocampal gyrus | 0.07 | 0.14 | 0.52 | 0.000000 |
| Supramarginal gyrus | 0.05 | 0.12 | 0.42 | 0.000000 |
| Caudal middle frontal gyrus | 0.06 | 0.16 | 0.37 | 0.000000 |
| Postcentral gyrus | 0.04 | 0.10 | 0.37 | 0.000000 |
| Temporal pole | 0.06 | 0.16 | 0.36 | 0.000000 |
| Isthmus of cingulate gyrus | 0.06 | 0.17 | 0.35 | 0.000000 |
| Superior temporal gyrus | 0.02 | 0.08 | 0.31 | 0.000000 |
| Lateral orbitofrontal cortex | 0.02 | 0.06 | 0.30 | 0.000000 |
| Fusiform gyrus | 0.03 | 0.11 | 0.30 | 0.000000 |
| Inferior temporal gyrus | 0.02 | 0.12 | 0.18 | 0.000000 |
| Superior frontal gyrus | 0.01 | 0.07 | 0.16 | 0.000000 |
| Lateral occipital cortex | 0.00 | 0.10 | 0.05 | 0.123153 |
| Medial orbitofrontal cortex | -0.01 | 0.12 | -0.06 | 0.040444 |
| Posterior cingulate gyrus | -0.01 | 0.15 | -0.09 | 0.002520 |
| Lingual gyrus | -0.01 | 0.11 | -0.10 | 0.001182 |
| Precentral gyrus | -0.01 | 0.07 | -0.14 | 0.000007 |
| Cuneus cortex | -0.03 | 0.14 | -0.23 | 0.000000 |
| Superior parietal cortex | -0.02 | 0.09 | -0.27 | 0.000000 |
| Rostral middle frontal gyrus | -0.05 | 0.08 | -0.63 | 0.000000 |
| Insula | -0.04 | 0.07 | -0.66 | 0.000000 |
| Precuneus cortex | -0.06 | 0.08 | -0.71 | 0.000000 |
| Paracentral sulcus | -0.13 | 0.14 | -0.89 | 0.000000 |
| Pericalcarine cortex | -0.11 | 0.12 | -0.90 | 0.000000 |
| Banks of superior temporal cortex | -0.25 | 0.24 | -1.04 | 0.000000 |
| Pars triangularis | -0.19 | 0.16 | -1.20 | 0.000000 |
| Middle temporal gyrus | -0.12 | 0.09 | -1.42 | 0.000000 |
| Pars orbitalis | -0.23 | 0.13 | -1.78 | 0.000000 |
| Frontal pole | -0.31 | 0.18 | -1.78 | 0.000000 |
| Inferior parietal cortex | -0.21 | 0.10 | -2.02 | 0.000000 |

**Fig. 4.**
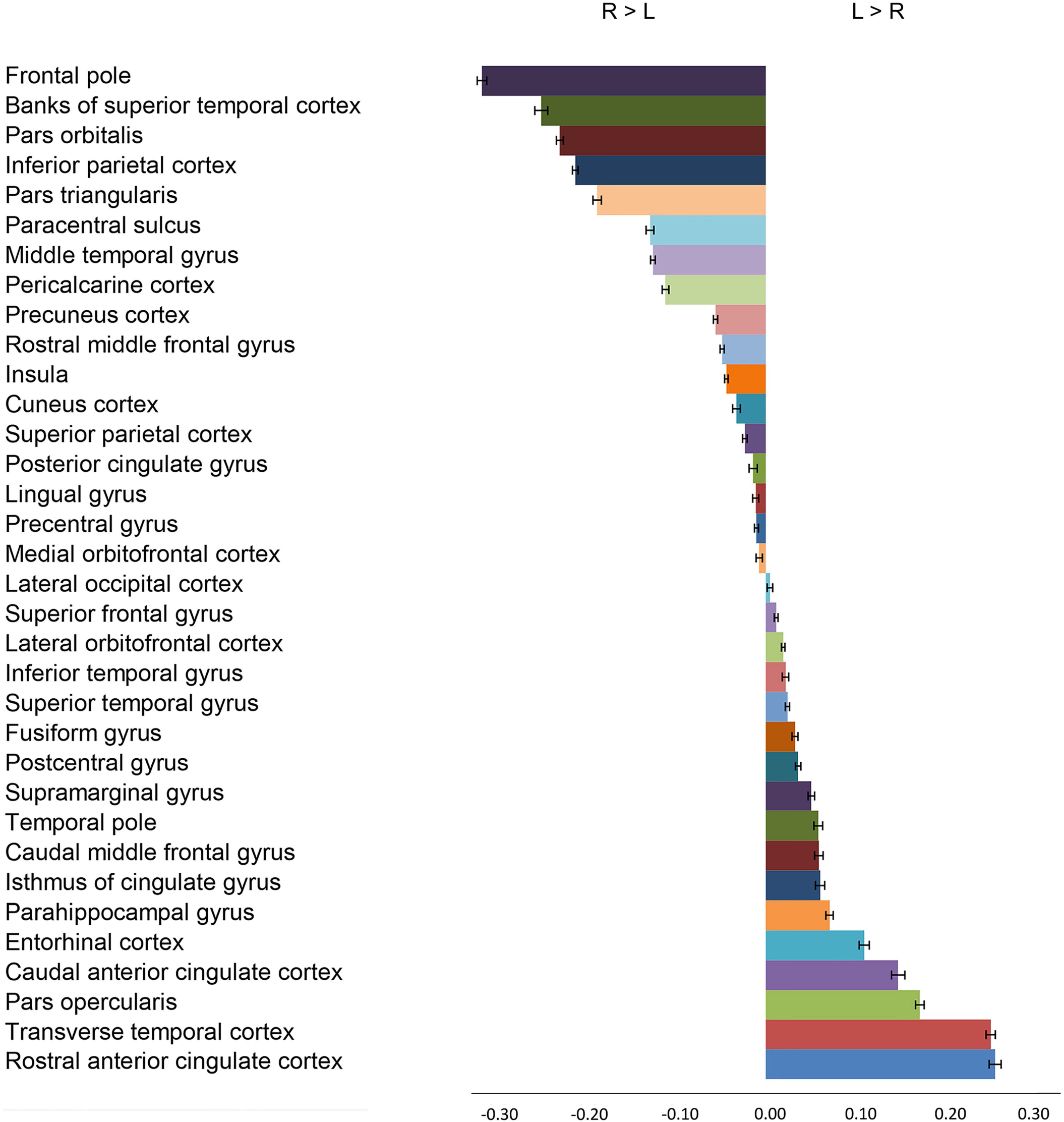
Asymmetry indices of gray matter volume in cortical regions. Positive asymmetry indicates leftward asymmetry (L > R), while negative asymmetry (R > L) indicates rightward asymmetry. Error bars represent SEM.

### Cerebellum, Hippocampus and Subcortical Structures

Mean of volumes of the cerebellar cortex and white matter, hippocampus, pallidum, putamen, caudate nucleus, nucleus accumbens and amygdala show a significant rightward asymmetry. Thalamus present a significant leftward asymmetry (Table.5 and Fig.5).

**Table 5.**
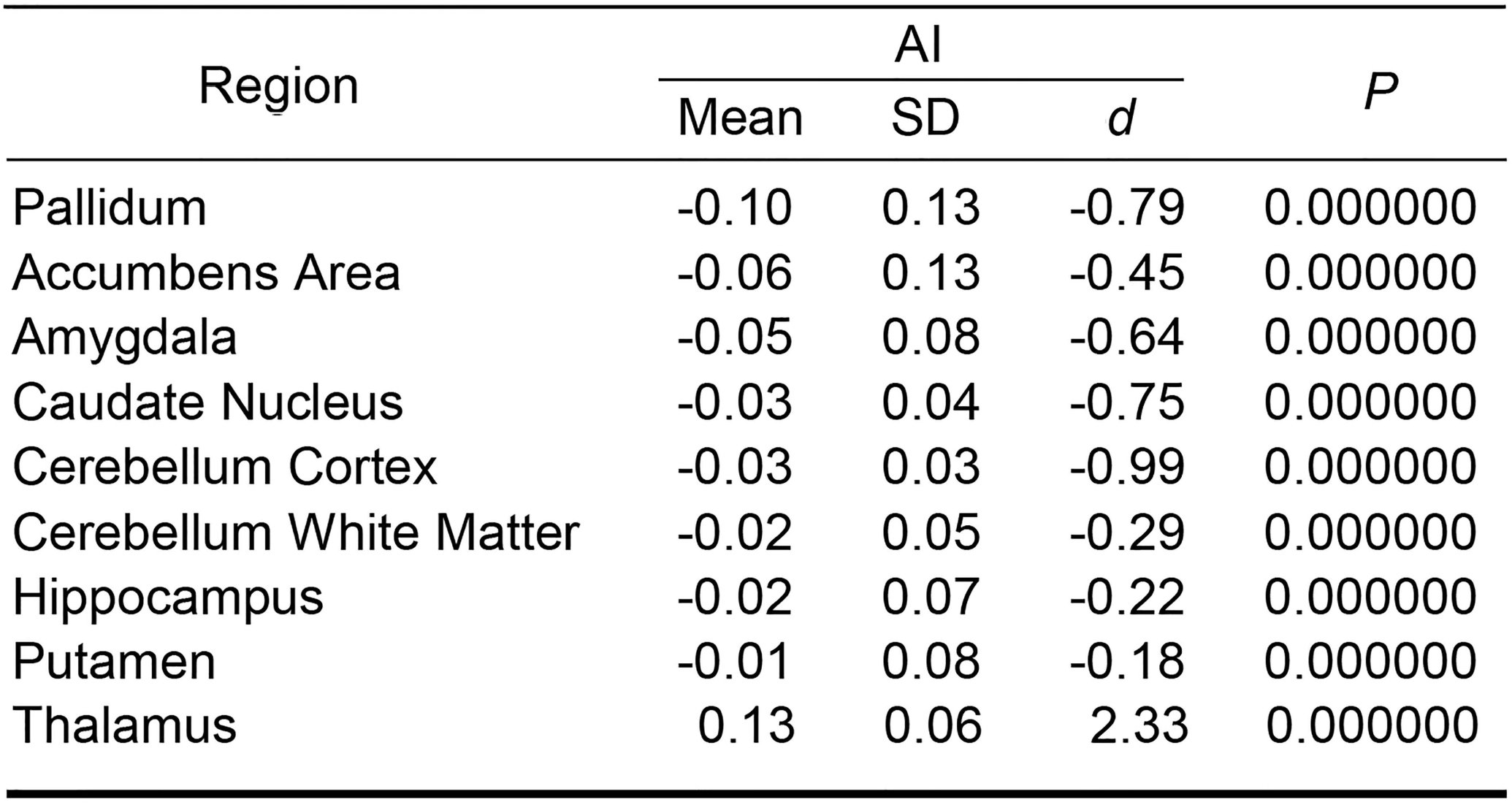
Asymmetry Index for hippocampus,cerebellum and subcortical structures

**Fig. 5.**
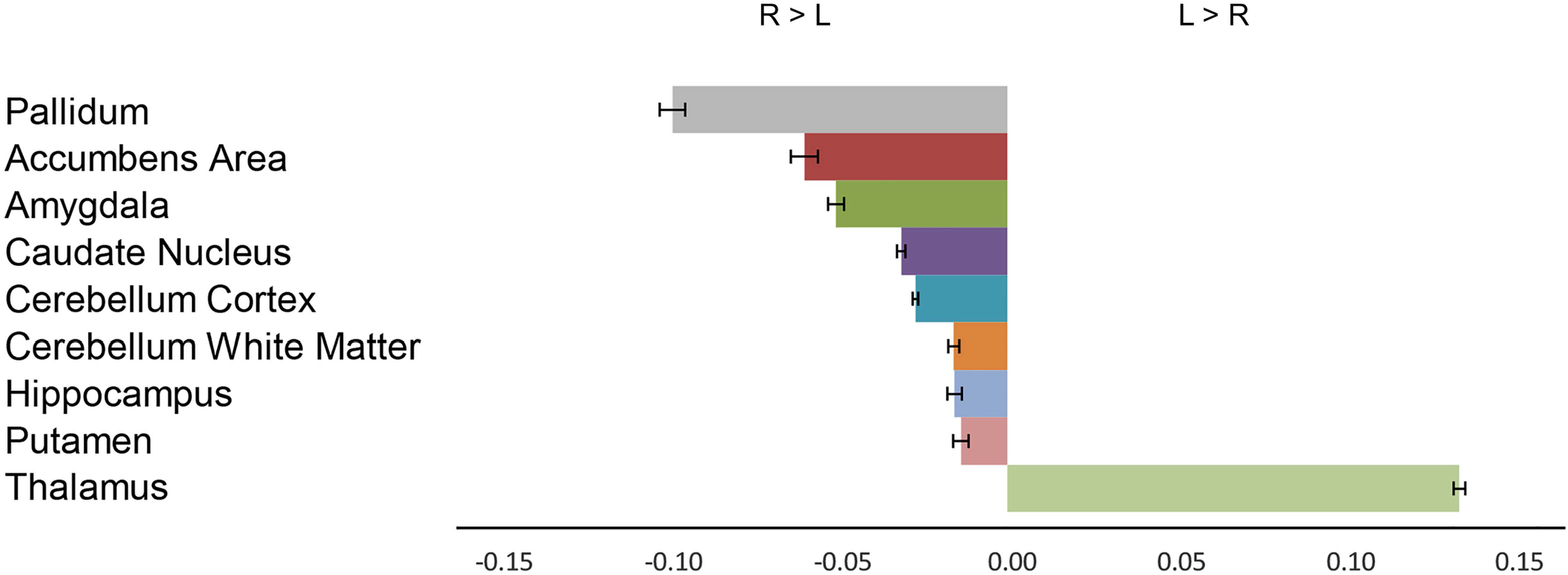
Asymmetry indices of hippocampus, cerebellum and subcotical regions. Positive asymmetry indicates leftward asymmetry (L > R), while negative asymmetry (R > L) indicates rightward asymmetry. Error bars represent SEM.

### Variation of the Cerebral Cortical Metrics

Cerebral cortical surface area (CV=14) and gray matter volume (CV=14) display more global variability than thickness (CV=6) (Fig.6A).

**Fig. 6.**
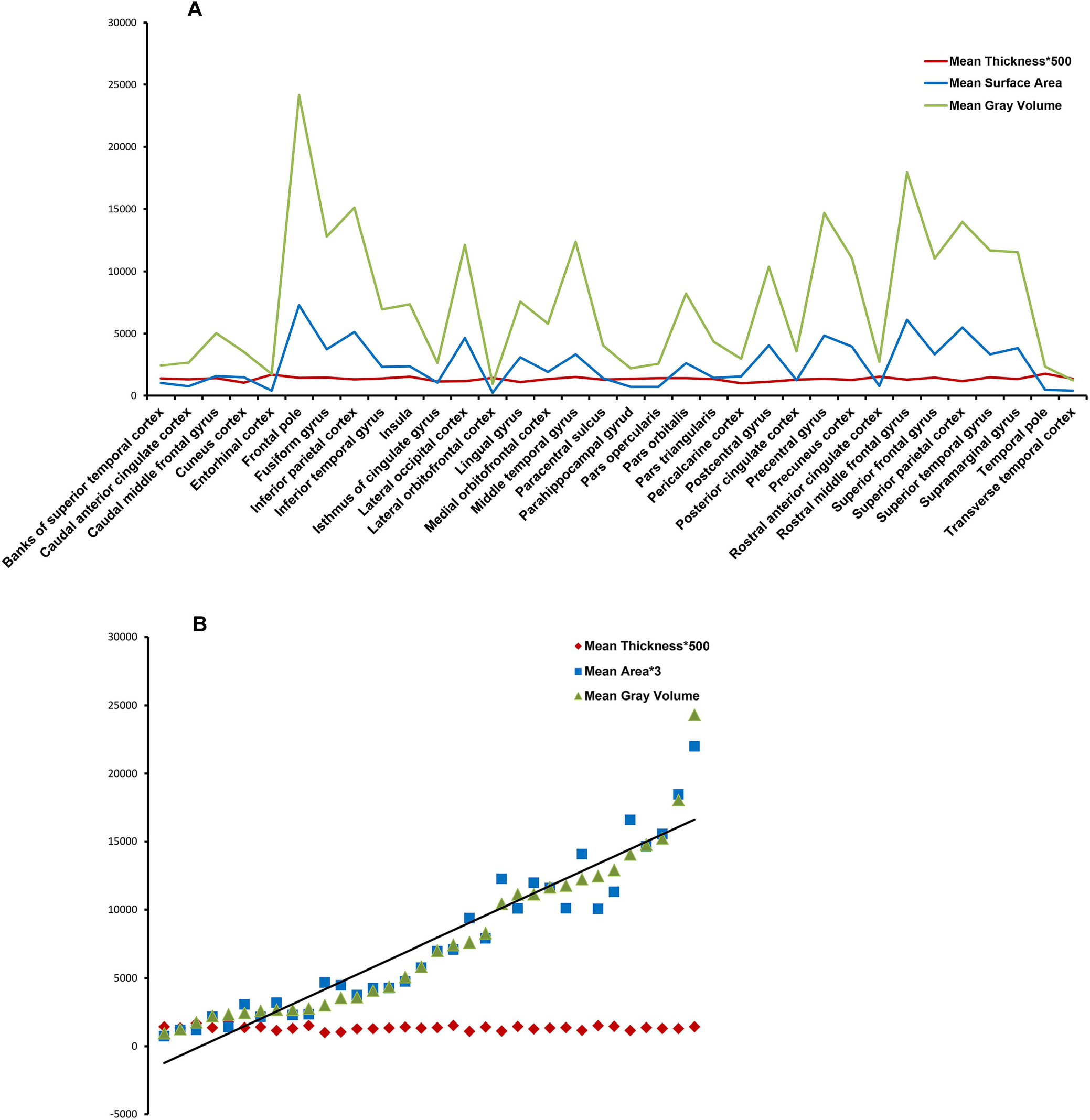
Cortical gray matter volume follows and strongly correlates with surface area across cortical regions (R2=0.96, p < 0000001) (A, B) However, cortical thickness does not show a significant correlation with surface area or gray matter volume (B). In addition, surface area (CV=14) and gray matter volume (CV=14) show more global variability compared with cortical thickness (CV=6) (A). Thickness and surface area values were multiplied by a constant to make comparison easier.

### Correlation between the Cerebral Cortical Metrics

We observed a tendency towards a negative correlation between cortical thickness and surface area across cortical regions (R^2^=0.04, *P* = 0.24) as well as the corresponding AIs (R^2^=0.1, *P* = 0.08) which do not reach the statistical significance. Furthermore, thickness and the gray matter volume do not show a significant correlation (R2=0.032, *P* = 0.31), neither their corresponding AIs (R2=0.003, *P* = 0.76406). However, surface area and gray matter volume (R^2^=0.96, *P* < 0000001) as well as their corresponding AIs (R^2^=0.6, *P* < 0000001) present a strong and significant correlation together (Fig.6 and 7).

### Gender

MANOVA revealed a significant effect of gender on AIs of the studied structures [F (113, 998) 1.9; *P* < 0. 0.0000005, partial η2=0.18]. However, even though, a number of nominally significant regions with exaggerated asymmetry patterns between genders was noted in *post-hoc* analysis, none survived Bonferroni correction (Table.6).

**Table 6.**
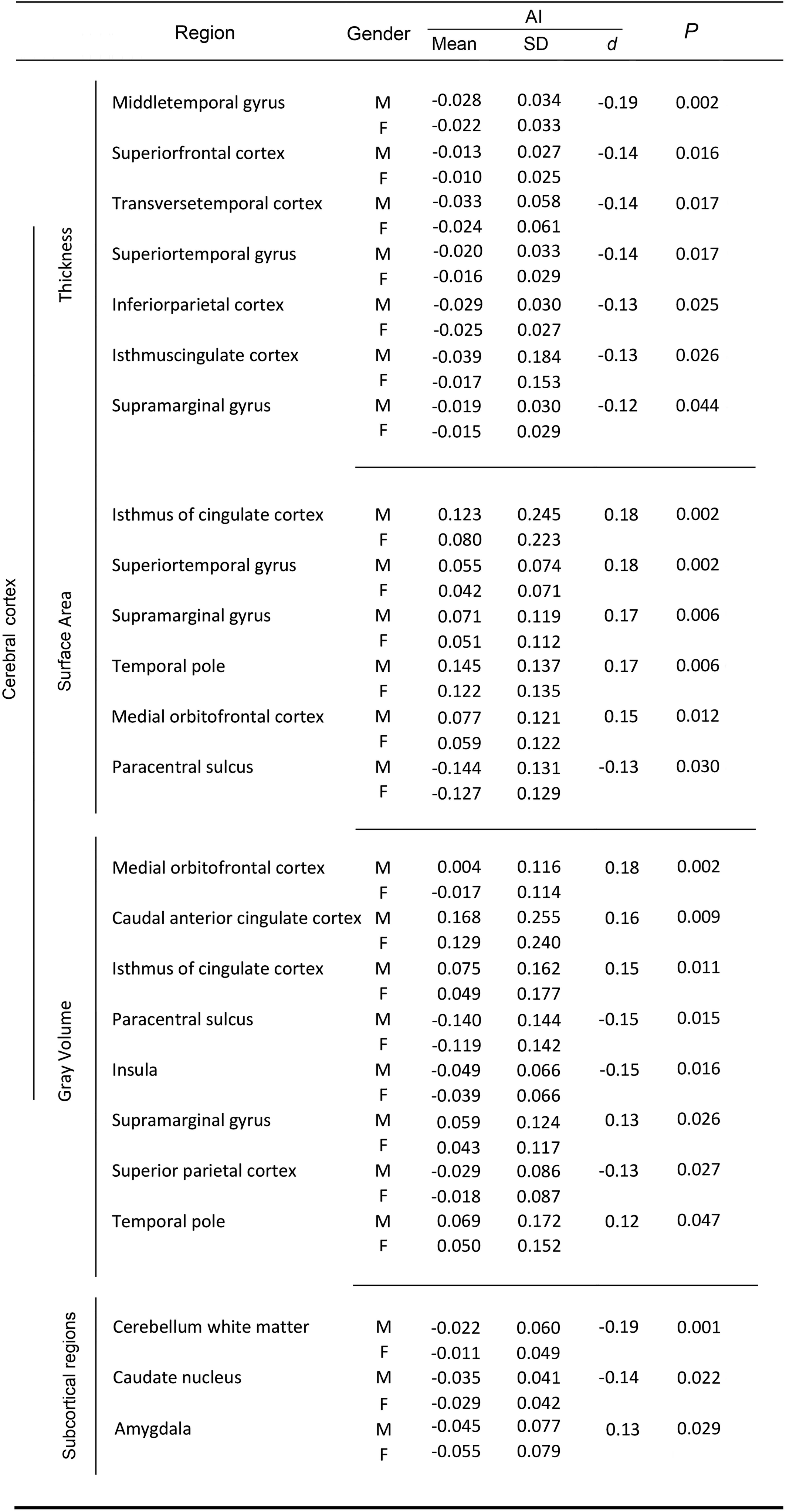
Regions with nominally signifiant gender effect on asymmetry index

### Age

Age does not show a significant effect on AIs of studied structures [F (113, 998) 1; *P* = 0.35, partial η2=0.11].

## DISCUSSION

To best of our knowledge, this is, by far, the largest study of asymmetry in healthy young adults which have been analyzed by standardized parcellation and imaging methods. Previous reports have either studied smaller groups which would not provide necessary statistical power or analyzed the data which have been obtained through heterogeneous methods. Moreover, this is the first study which has studied asymmetry in three metrics of the cerebral cortex as well as cerebral white matter, cerebellar cortex and white matter, hippocampus and six subcortical structures in the same group of individuals. A major finding of this study is the almost ubiquitous rightward asymmetry in mean of gray and white matter volumes in the studied structures. Volumes of the cerebral as well as cerebellar cortex and white matter, hippocampus and all subcortical structures expect thalamus show a significant rightward asymmetry. In addition, mean global cerebral cortical thickness and surface area present a rightward asymmetry as well (Table.1 and 5), (Fig.1 and 5). The findings are in agreement with a previous study describing a rightward asymmetry in the cortical gray matter as well as cerebral white matter volumes (24). In addition, although, only a few studies has addressed hemispheric asymmetry in cortical surface area, thickness and gray matter volume in the same group of participants, a global rightward asymmetry in all these metrics has been reported before (25). Moreover, rightward asymmetry of the cerebral gray matter volume, and cortical surface area has been described in other studies (13, 26). Nevertheless, conflicting asymmetry patterns have been also reported (12, 13, 27, 28). Despite the global rightward asymmetry of cortical metrics, differential patterns at regional level were observed. Although, most cortical regions show a rightward asymmetry in thickness (Table.2 and Fig.2), surface area presents a more even distribution between two hemispheres with 53% of regions showing a significant leftward asymmetry (Table.3 and Fig.3). It is notable that in the lateral surface of the cerebral cortex, asymmetry in surface area shows a relationship with the distance to Sylvian fissure leading to a leftward asymmetry in peri-Sylvian regions which has been frequently reported before (13, 25, 29–31). For example, both caudal middle and inferior frontal (pars opercularis) gyri display a leftward asymmetry in surface area. However, rostral middle and inferior frontal (pars orbitalis and pars triangularis) gyri show a rightward asymmetry. A similar pattern is observed in parietal cortex. Postcentral and supramarginal gyri show a leftward asymmetry in surface area. However, inferior and superior parietal cortices show a rightward asymmetry. In medial surface of cerebrum an anterior-left, posterior-right pattern of asymmetry in surface area is observed. Superior frontal as wells as rostral anterior cingulate regions show a leftward asymmetry. However, precuneous, cuneous, caudal anterior and posterior cingulate cortices show a rightward asymmetry. Between structural variables of cortical gray matter volume (32), surface area shows more global variability (CV=14) compared with thickness (CV= 6) (Fig.6A), which is in agreement with previous reports (29, 33). This is probably why global variability (CV=14%) as well as the volume of cortical gray matter often follows cortical surface area (R^2^=0.96, *P* < 0000001) (Fig.6). Concordantly, the asymmetry patterns of the cortical gray matter volume mostly pursue surface area (Fig.7A). This includes leftward asymmetry of cortical gray matter volume in peri-Sylvian regions including transvers temporal cortex, superior temporal gyrus, temporale pole, pars opercularis and supramarginal gyrus which is in agreement with previous reports (Table.4 and Fig.4) (9, 10, 26, 34). Regions with concordant rightward asymmetry between surface area and gray matter volume include middle temporal gyrus, pars orbitalis, pars triangularis, insula, inferior parietal, cuneous, precuneous and posterior cingulate cortices which are mostly in line with a previous study (30). Regions with opposite asymmetry pattern between surface area and gray matter volume are not numerous and include banks of superior temporal sulcus, caudal rostral cingulate and medial orbitofrontal gyri where gray matter volume follow asymmetry pattern of the cortical thickness (Table.2, 4 and Fig.2, 4). Consistently, unlike a previous report (25), we noticed a strong global relationship between AIs of the cortical surface area and gray matter (R=0.6, *P* < 0000001) compared with a non-significant relationship between AIs of the cortical thickness and gray matter (R2=0.003, *P* = 0.76406) which is in line with a previous report (33). Fig. 7B). This supports the notion of independent character of cortical thickness and surface area as neuroanatomical unites (30, 33, 35). A good example of how variation in surface area influences asymmetry of the gray matter volume is Broca’s area. Although, both parts of Broca’s area, pars triangularis and pars opercularis, present a rightward asymmetry in thickness, the gray matter volume follows asymmetry pattern of surface area, i.e. rightward in pars triangularis and leftward in pars opercularis. In fact, difference in surface area asymmetry places the gray matter volume asymmetry of pars triangularis (*d*=−1.2) and pars opercularis (*d*=1.1) among strongest opposite asymmetries in whole brain. Interestingly, a similar pattern is observed in another language related region, supramarginal gyrus i.e. a rightward asymmetry in thickness and leftward asymmetry in surface area and gray matter volume. Although, many of the asymmetry patterns in cortical metrics described here, has been reported in different studies (9, 13, 25, 26, 28–30), it is the first time, in our knowledge, that above mentioned relationship between cortical metrics in determining asymmetry patterns across cerebral cortex has been described. The results of this study in regional cortical asymmetry are similar to a previous report (30). However, probably lower number of cases and lack of necessary statistical power in addition of strict correcting for multiple comparisons by Bonferroni procedure has led to many statistically insignificant results in above mentioned study. In addition, we could replicate many of the asymmetry patterns reported by other studies (12, 29, 31), even though, inconsistencies exist which detailed discussion about it is beyond the scope of this paper. However, we have to emphasize that a part of the inconsistencies could be due to the age of participants as reports suggest it affects asymmetry patterns (17, 18). On the other hand, although, our results show similarities with another large data analysis (13), especially on asymmetries in cortical surface area, we could not replicate reported leftward hemispheric asymmetry as well as many regional asymmetries in cortical thickness. Important heterogeneity in asymmetry results between analyzed data sets which has been acknowledged by the authors could be one of the reasons of this discrepancy (13).

**Fig. 7.**
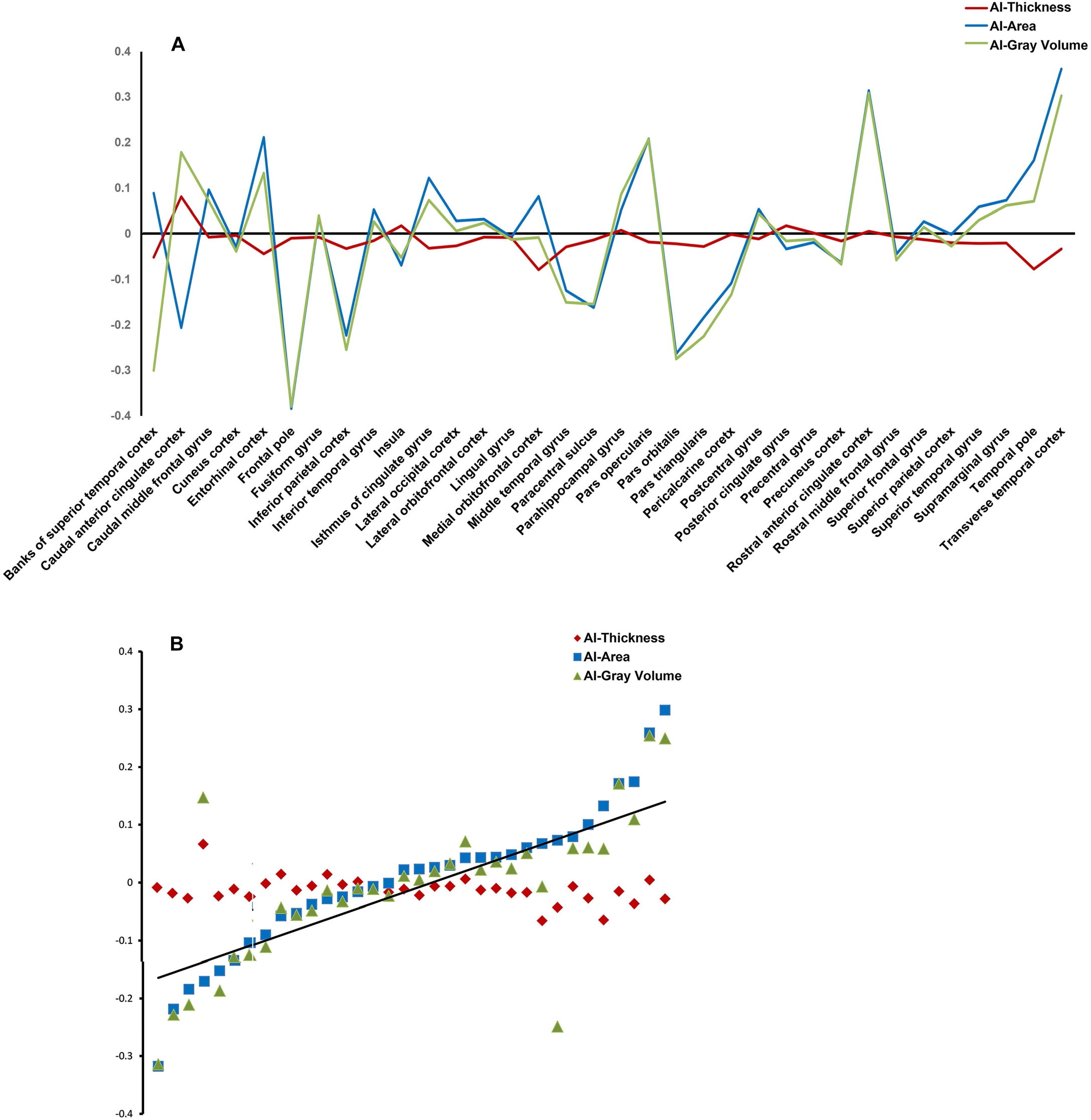
Asymmetry indices of cortical gray matter volume frequently follow and strongly correlate with asymmetry indices of surface area (R2=0.6, p < 0000001) (A, B). However, asymmetry indices of cortical thickness do not show a significant correlation with asymmetry indices of surface area or gray matter volume (B).

### Cerebellum, Hippocampus and Subcortical Structures

Although, fewer studies on the asymmetry of subcortical structures are available, discrepancies prevail here as well. Apart from putamen and pallidum, the observed rightward asymmetry in hippocampus, amygdala, caudate nucleus and nucleus accumbens as well as leftward asymmetry in thalamus are in agreement with the largest met-analysis published on asymmetry of subcortical structures (11) as well as other reports (27, 36). The reason of the reported leftward asymmetry in putamen and pallidum in above mentioned studies can be a more rapid decline in volume of right putamen and pallidum and a leftward shift in the asymmetry with age which has been reported before (11, 37). The observed rightward asymmetry in the cerebellum is in agreement with previous studies as well (26, 27, 38). Although, a leftward asymmetry has been also reported (10). Regarding the effects of gender on brain asymmetries, while, no region survives Bonferroni correction in *post hoc* analysis, a few points about observed nominally significant asymmetries (Table.6) worth to be discussed. First, about 12% of cortical regions show a nominal gender effect on at least one of the metrics. Second, most of exaggerated tendencies to either side belong to male brain. This suggests male probably has a more lateralized brain which is in agreement with previously reported structural as well as functional studies (1, 10, 27, 39, 40). Third, the observed asymmetries are relatively subtle, with a Cohen’s-*d* range of −0.19 to 0.19 (Table.6) which is also in line with previous reports (27, 41). Fourth, an important part of cortical language as well as visuospatial regions display following gender specific tendencies in male. Transverse temporal cortex, superior temporal and supramarginal gyri as well as inferior parietal cortex present an exaggerated rightward asymmetry in thickness. In addition, surface area displays an exaggerated leftward asymmetry in temporal pole, superior temporal and supramarginal gyri. Finally, an exaggerated leftward asymmetry in gray matter volume of temporal pole and supramarginal gyrus as well as a rightward asymmetry in superior parietal cortex was noted. The observed accentuated asymmetries in male are in agreement with reports indicating more left lateralized language (39) and right lateralized visuospatial functions in males (42). Furthermore, medial orbitofrontal cortex gray matter volume present an opposite tendency in asymmetry between genders i.e. rightward in female and leftward in male (*P* < 002, uncorrected). In addition, amygdala present an exaggerated rightward asymmetry in female. Although, several lines of evidence suggest that medial prefrontal cortex and amygdala show a differential asymmetrical organization between genders which contribute to distinct cognitive and affective information processing, underlying mechanisms are far from being clear (43–46). The presence (10, 13, 14, 47) or absence of a significant effect of gender on asymmetry (18, 27, 28, 31, 34, 48) has been frequently reported. In addition of previous mentioned reasons of inconsistencies, this could reflect the fact that asymmetry as a principal developmental feature of human brain follows similar trajectories in both genders yielding only to subtle differences (18, 41). In addition, some reports suggest that gender effects on asymmetry might change with age (17). Furthermore, since the absolute values of intracranial volume as well as body size are larger in males, it is challenging to differentiate gender effects on asymmetry from those related to above mentioned factors (13, 31, 49). Finally, we did not observed a significant age effects on the asymmetry indices of structures studied. However, this obviously would not preclude a relationship between age and asymmetry as participants in this project has been chosen from young adults. This study, for the first time, has provided the asymmetry picture of whole brain structures in unprecedented number of carefully chosen young adults yielding to highly significant results. This provides a basis which would help to understand better this important hallmark of human brain.

## Acknowledgements

The project was supported by funds from Shahid Behehsti University. Data were provided by the Human Connectome Project, WU-Minn Consortium (Principal Investigators: David Van Essen and Kamil Ugurbil; 1U54MH091657) funded by the 16 NIH Institutes and Centers that support the NIH Blueprint for Neuroscience Research; and by the McDonnell Center for Systems Neuroscience at Washington University.

